## Extended Data Figures 1 - 11 for "Vesicular release probability sets the strength of individual Schaffer collateral synapses"

ED Figure 1: Temporal stability of iGluSnFR transients.

ED Figure 2: Measured release probabilities are independent of imaging noise.

ED Figure 3: Two sigma threshold to discriminate successes from failures.

ED Figure 4: Multisynaptic boutons show a second release site under high release probability.

ED Figure 5: Low occupancy of AMPARs.

ED Figure 6: Unchanged synaptic release probability upon block of NMDAR.

ED Figure 7: Expression of iGluSnFR does not affect synaptic transmission.

ED Figure 8: Histogram bin size effects.

ED Figure 9: Extracted parameters are robust to assumed saturation value.

ED Figure 10: Correlation matrix of extracted quantal parameters.

ED Figure 11: Extracting iGluSnFR kinetics from synaptic imaging data.

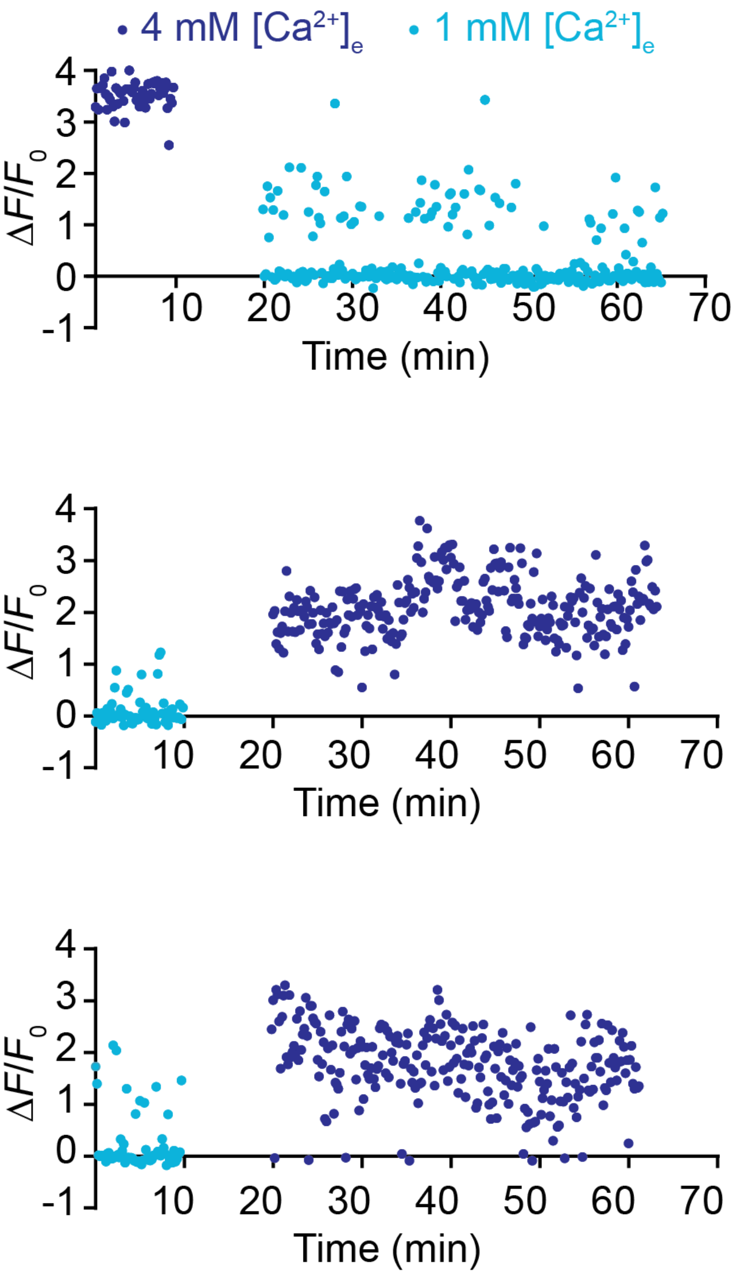


**Extended Data Figure 1: Temporal stability of iGluSnFR transients.**

Response amplitude in response to single presynaptic APs monitored in ACSF containing 1 and 4 mM $[Ca^{2+}]_e$ at 33°C. During the wash-in period (10 min) from 1 to 4 mM $[Ca^{2+}]_e$ or vice versa, no images were acquired. Three different Schaffer collateral boutons from 3 different organotypic cultures.

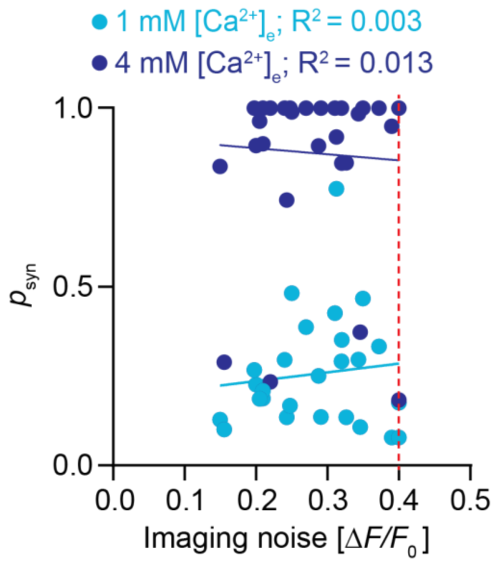


**Extended Data Figure 2: Estimated release probabilities were independent of imaging noise.** $p_{syn}$ plotted as a function of the imaging noise. The imaging noise corresponds to the full width at half maximum (FWHM) of a Gaussian fit to the distribution of the baseline noise from trial to trial for a given boutons. Experiments with an imaging noise above a FWHM of 0.4 $\Delta F/F_0$ (red dotted line) were discarded.

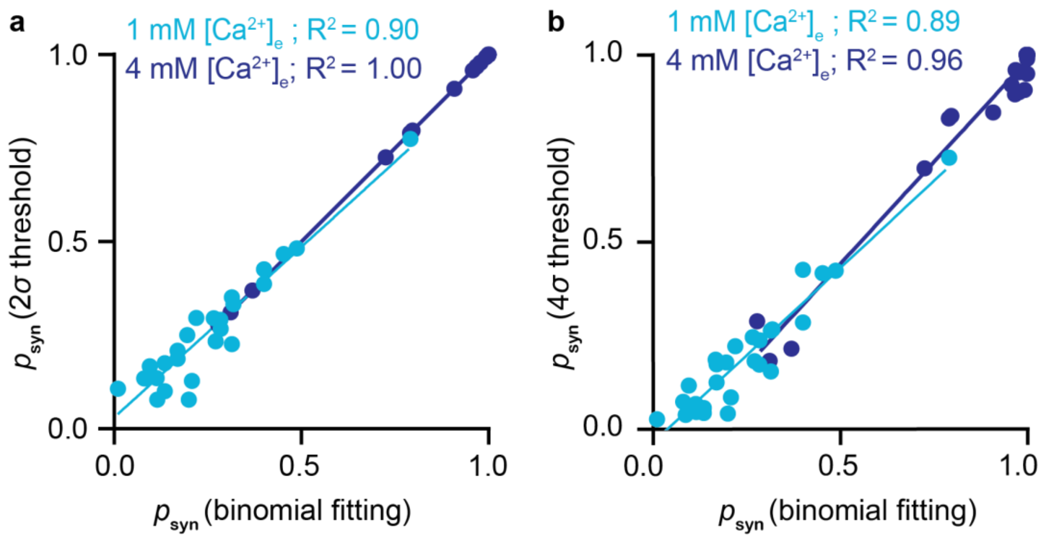


**Extended Data Figure 3: Two sigma threshold to discriminate successes from failures delivers $p_{syn}$ values consistent with full quantal analysis.**

**a)** Synaptic release probability ($p_{syn}$) measured as the number of successes (responses exceeding 2$\sigma$ of baseline fluorescence fluctuations) divided by the number of trials, plotted as a function of $p_{syn}$ calculated from the quantal parameters extracted with the binomial fitting procedure ($p_{syn} = 1-(1-p_{ves})^N$). Linear fits show high correlation between the two estimates of $p_{syn}$. **b)** Same as **a**, but classified with a higher threshold (4σ). The resulting estimates of $p_{syn}$ are generally lower and not as consistent with our binomial quantal analysis.

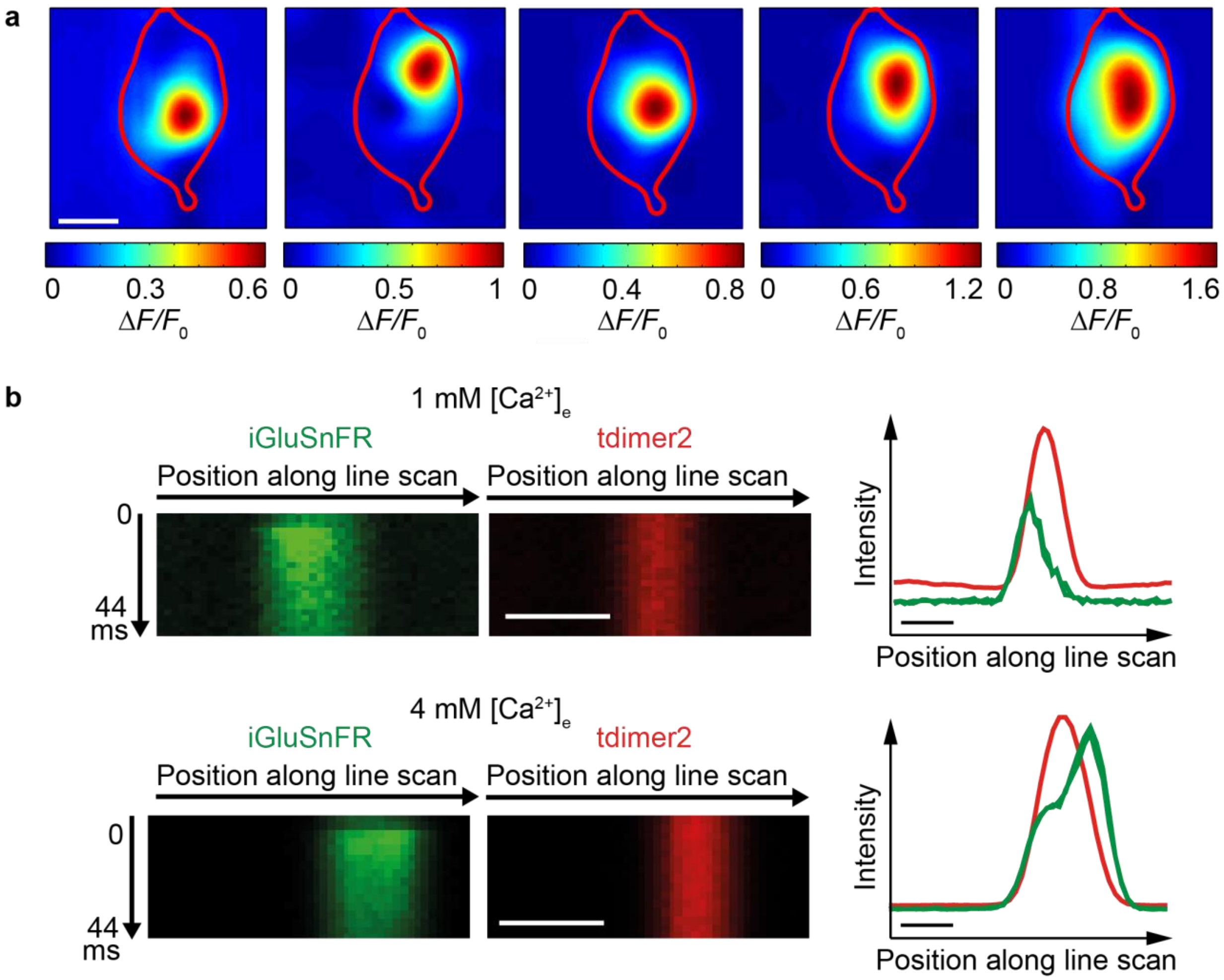


**Extended Data Figure 4: Multisynaptic boutons show a second release site under high release probability conditions. a)** Example of color-coded iGluSnFR signal of a multi-synapse bouton. Each frame represents the iGluSnFR signal in response to a single action potential. Scale bar, 1 µm. **b)** Example of the average intensity profile (average of ~ 30 trials) *post hoc* aligned on the red morphological marker of a straight line scan across a multisynaptic bouton in 1 mM $[Ca^{2+}]_e$ (*upper panel*) with a single hot spot and in 4 mM $[Ca^{2+}]_e$ (*lower panel*) with the appearance of a second hot spot. Boutons with multiple release sites were rare; we excluded them from further analysis. Scale bar, 1 µm.

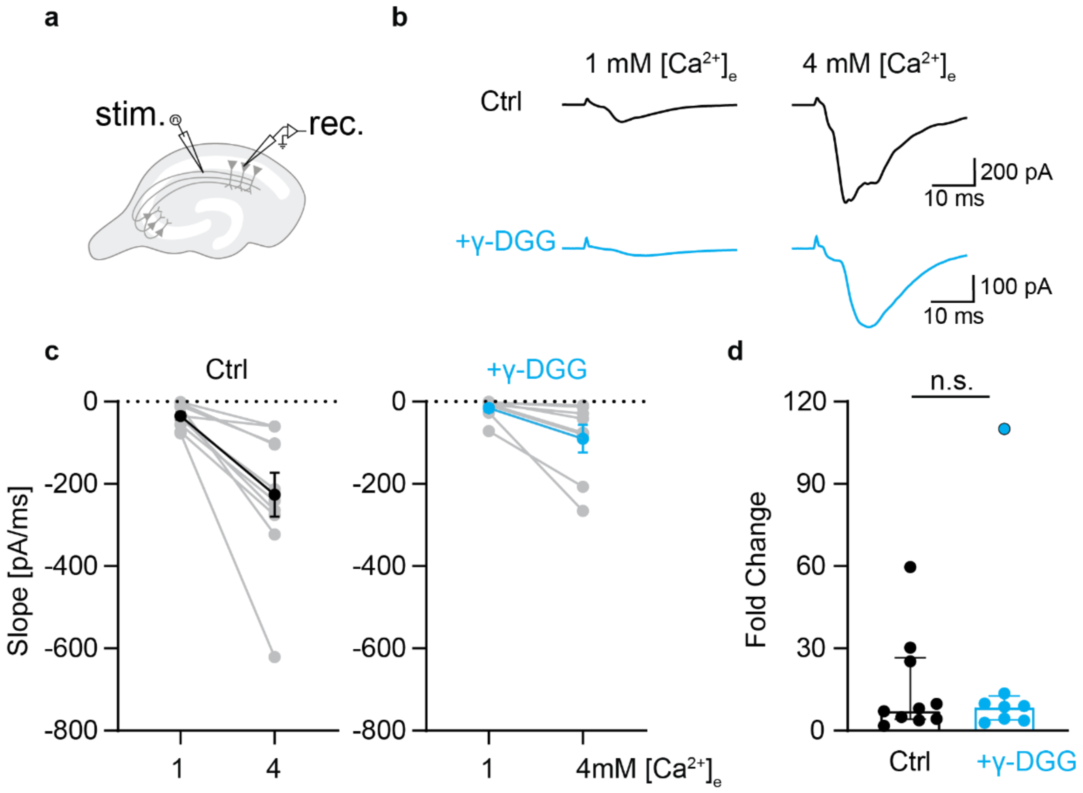


**Extended Data Figure 5: Low occupancy of AMPARs. a)** A monopolar electrode placed in *stratum radiatum* was used to elicit postsynaptic responses measured in whole cell voltage clamp from a CA1 neuron located ~300 µm away. To isolate AMPARs, the NMDAR blocker CPPene (10 µM) was added to the bath. **b)** Example traces of AMPARs responses measured in response to electrical field stimulation in control condition *(upper traces)* and in the presence of 10 mM γ-DGG, a competitive AMPARs antagonist *(lower traces)* which led to decreased evoked EPSCs. **c)** *(Left panel)* Slope measurements of evoked EPSCs in control condition in 1 mM $[Ca^{2+}]_e$ : -35.1 ± 8.8 pA/ms and in 4 mM $[Ca^{2+}]_e$: -226.5 ± 53.2 pA/ms, $n$ = 10 cells. *(Right panel)* Slope measurements upon application of γ-DGG in 1 mM $[Ca^{2+}]_e$: -15.5 ± 8.5 pA/ms and in 4 mM $[Ca^{2+}]_e$: -90.1 ± 33.7 pA/ms, $n$ = 8 cells. Values are plotted as mean ± SEM. **d)** Decrease of AMPARs occupancy did not lead to a significant increase in postsynaptic fold change from low to high $[Ca^{2+}]_e$. There is no significant fold change from 1 mM to 4 mM $[Ca^{2+}]_e$ in the absence (median: 7.57 fold, IQR: 4.1–26.5 fold, $n$ = 10 cells) or presence of γ-DGG (median: 8.84 fold, IQR: 3.8–12.6 fold, $n$ = 8 cells) (Mann Whitney test, $p$ = 0.97). Values are plotted as median with IQR.

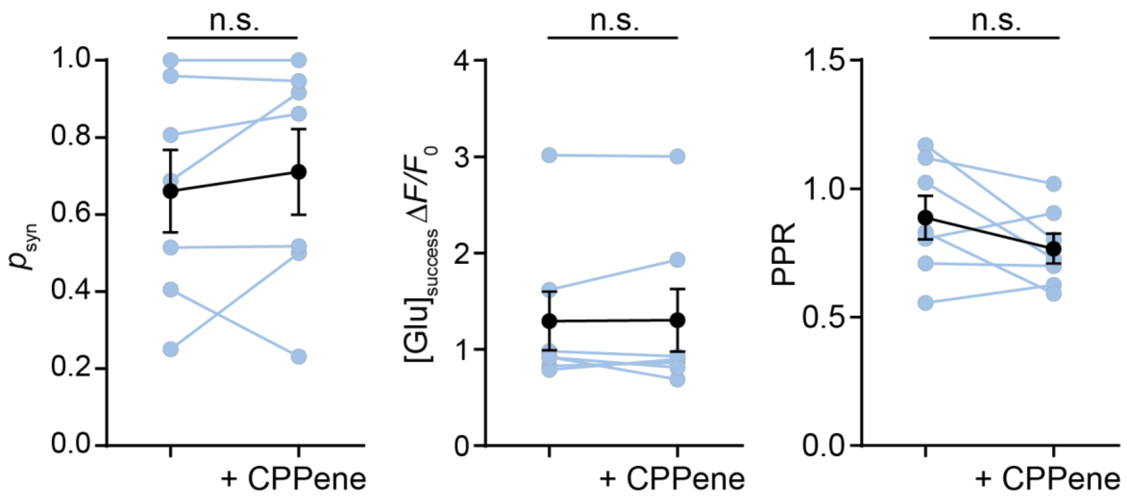


**Extended Data Figure 6: Unchanged synaptic release probability upon block of NMDAR.** iGluSnFR changes in fluorescence (single bouton, ~30 consecutive trials) were measured in response to single action potentials before and after the application of the NMDARs blocker CPPene (10 µM). There was no significant difference under NMDARs block in synaptic release probability, Ctrl: 0.66 ± 0.1, NMDAR block: 0.71 ± 0.1, paired t-test, $p$ = 0.41; cleft glutamate, Ctrl: 1.3 ± 0.3 $\Delta F/F_0$, NMDAR block: 1.3 ± 0.3 $\Delta F/F_0$, Wilcoxon test, $p$ = 0.94; and PPR, Ctrl: 89% ± 9%, NMDAR block: 77% ± 6%, paired t-test, $p$ = 0.13; $n$ = 7 boutons in 7 slices. Values are plotted as mean ± SEM.

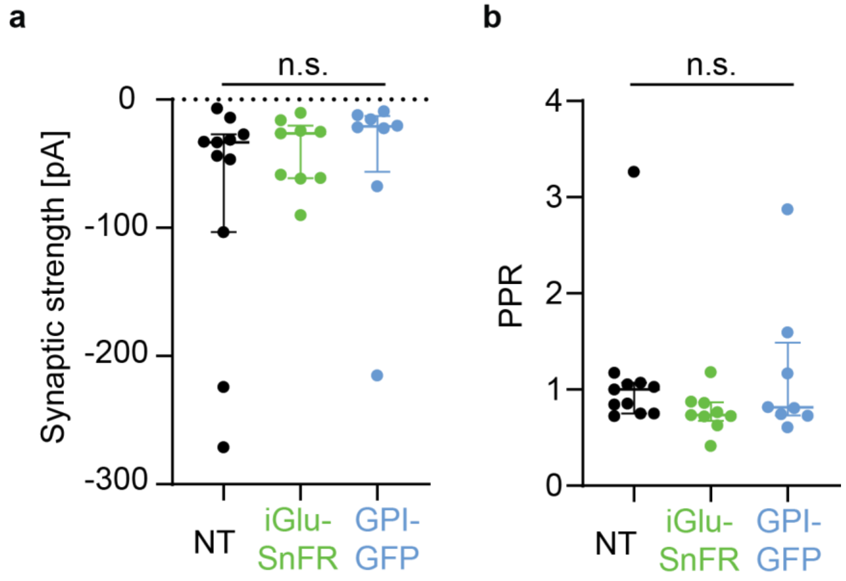


**Extended Data Figure 7: Expression of iGluSnFR does not affect synaptic transmission. a)** EPSCs were measured by dual patch-clamp recordings from connected CA3-CA1 pyramidal cell pairs in 4 mM $[Ca^{2+}]_e$. The NMDA blocker CPPene (10 µM) was added to the bath to isolate AMPAR responses. EPSCs were recorded from non-transfected (NT) CA3-CA1 pairs (median synaptic strength: -33.4 pA, IQR: -103.3 – -27.2 pA, $n$ = 11 pairs); iGluSnFR-expressing CA3-CA1 pairs (median synaptic strength: -26.4 pA, IQR: -61.4 – -20.25 pA, $n$ = 9 pairs); GPI-GFP-expressing CA3-CA1 pairs (median synaptic strength: -21.0 pA, IQR: -56.2 – -12.9 pA, $n$ = 8 pairs). There was no significant difference in measured synaptic strength between the different groups of connected CA3-CA1 pairs (Kruskal-Wallis test, $p$ = 0.31). **b)** EPSCs showed paired-pulse depression (ISI = 48 ms) in 4 mM $[Ca^{2+}]_e$ for NT CA3-CA1 pairs (median PPR: 100%, IQR: 75% –107%, $n$ = 11 pairs), for the iGluSnFR expressing CA3-CA1 (median PPR: 74%, IQR: 68% – 87%, $n$ = 9 pairs) and for the GPI-GFP CA3 expressing-CA1 pairs (median PPR: 81%, IQR: 73%– 149%, $n$ = 8 pairs). There was no significant difference in measured synaptic strength between the different groups of connected CA3-CA1 pairs (Kruskal-Wallis test, $p$ = 0.17). Values are plotted as median with IQR.

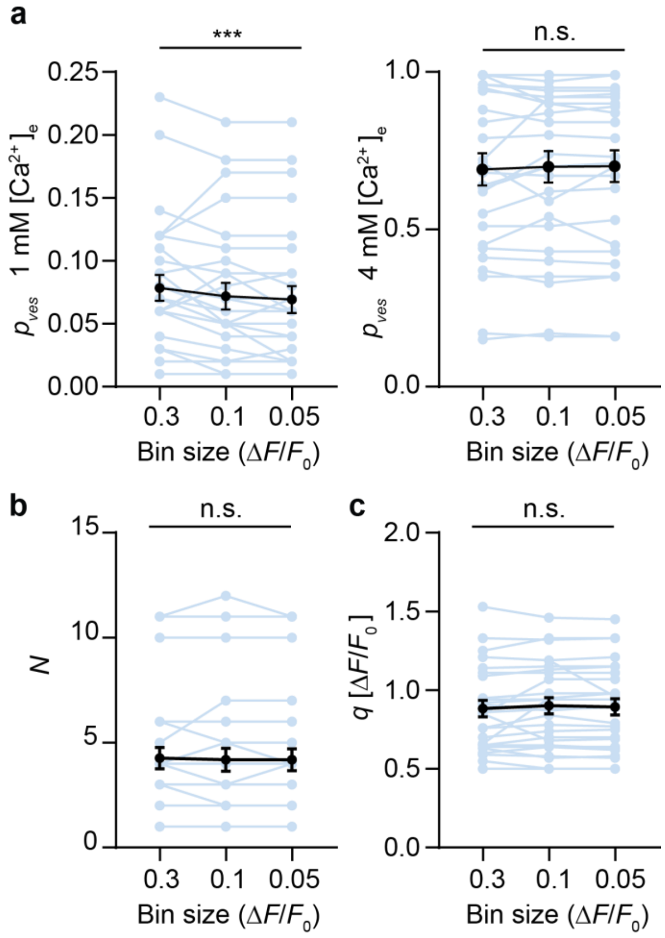


**Extended Data Figure 8: Quantal analysis by fitting binomial distributions to histograms is robust with respect to histogram bin size.**

**a)** $p_{ves}$ in 1 mM $[Ca^{2+}]_e$ (*left*) and 4 mM $[Ca^{2+}]_e$ (*right*) extracted by binomial fitting of response amplitude histograms for different bin sizes. Mean $p_{ves}$ in 1 mM $[Ca^{2+}]_e$ was different for bin size 0.3 $\Delta F/F_0$ (0.08 ± 0.01), bin size 0.1 $\Delta F/F_0$ (0.07 ± 0.01) and bin size 0.05 $\Delta F/F_0$ (0.07 ± 0.01) (Friedman test, $p$ = 0.0007, $n$ = 27 boutons), but the size of the effect was small. Mean $p_{ves}$ in 4 mM $[Ca^{2+}]_e$ did not depend on bin size (one-way ANOVA, $p$ = 0.5, $n$ = 27 boutons). **b)** Number of vesicles $N$ extracted by binomial fitting did not depend on bin size (Friedman test, $p$ = 0.4, $n$ = 27 boutons). **c)** Quantal size $q$ extracted by binomial fitting did not depend on bin size (one-way ANOVA, $p$ = 0.45, $n$ = 27 boutons). Values are plotted as mean ± SEM.

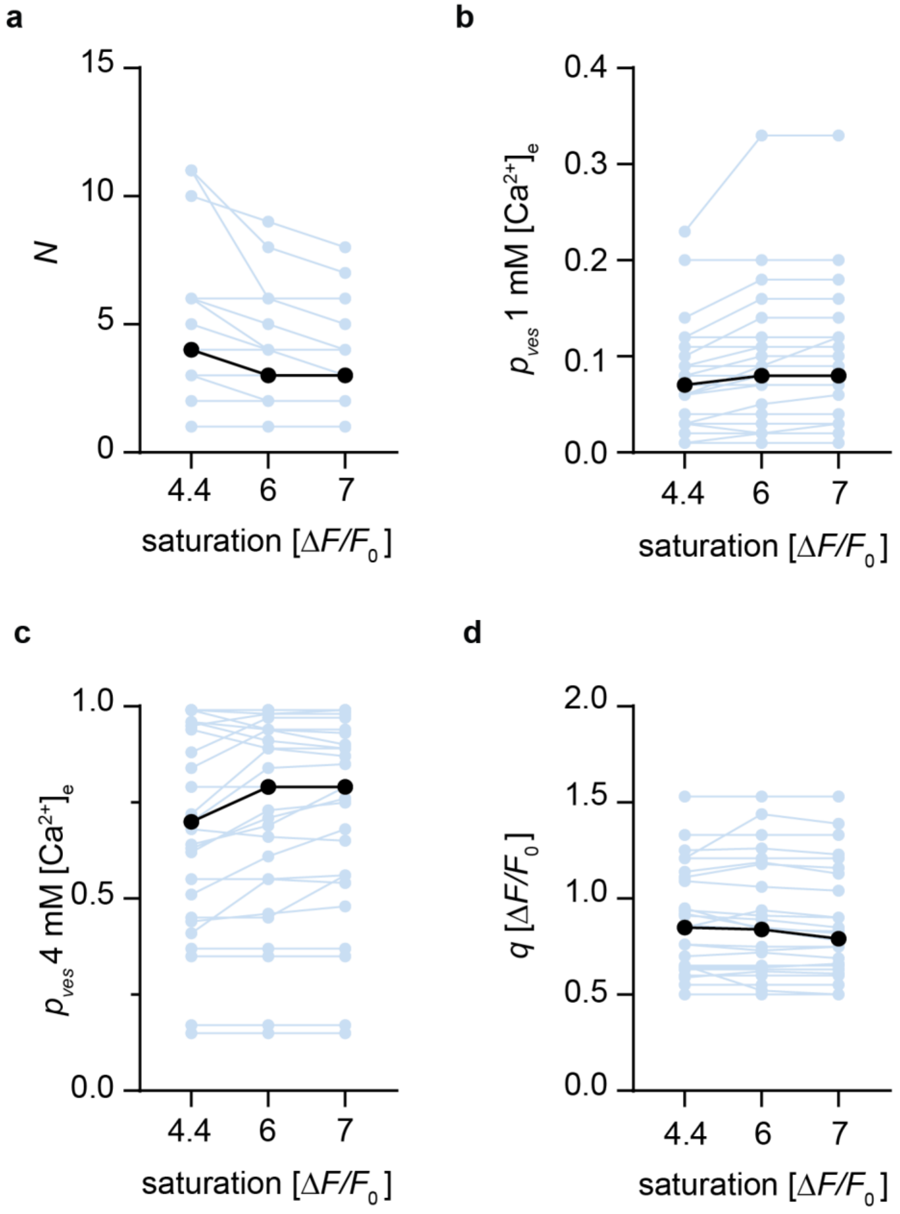


**Extended Data Figure 9: Extracted parameters are robust to the assumed saturation value.** Extracted number of **a)** vesicles **b)** vesicular release probability in 1 mM [$Ca^{2+}$]$_e$ **c)** vesicular release probability in 4 mM [$Ca^{2+}$]$_e$ and **d)** quantal size from the binomial fitting procedure of individual boutons assuming different saturation values of iGluSnFR-expressing boutons ($n$ = 27 boutons). Median values are plotted in black. Our Monte-Carlo simulations (Fig. 5f) suggest apparent saturation at 6 ΔF/F for a synapse at optimal orientation and 7 ΔF/F for a 40° tilted synapse. Assuming saturation at 4.4, which is the literature value of iGluSnFR when calibrated with glutamate-containing solutions, requires one more vesicle (on average) to fit the shape of the response histograms.

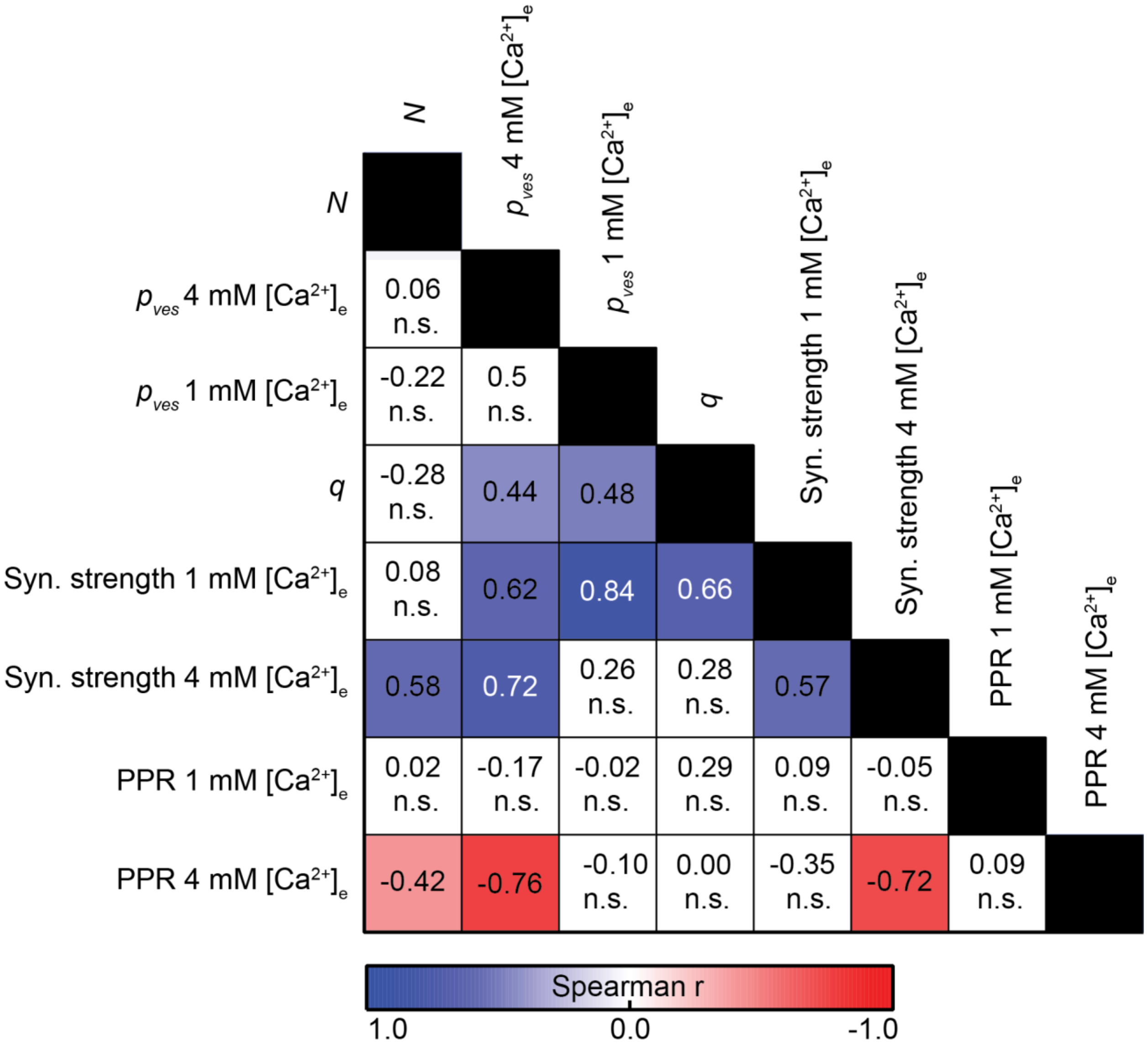


**Extended Data Figure 10: Correlation matrix of extracted quantal parameters, synaptic strength (*p**q*N*) and paired-pulse ratio (PPR).** Significant positive correlations are labeled in blue, significant negative correlations in red.

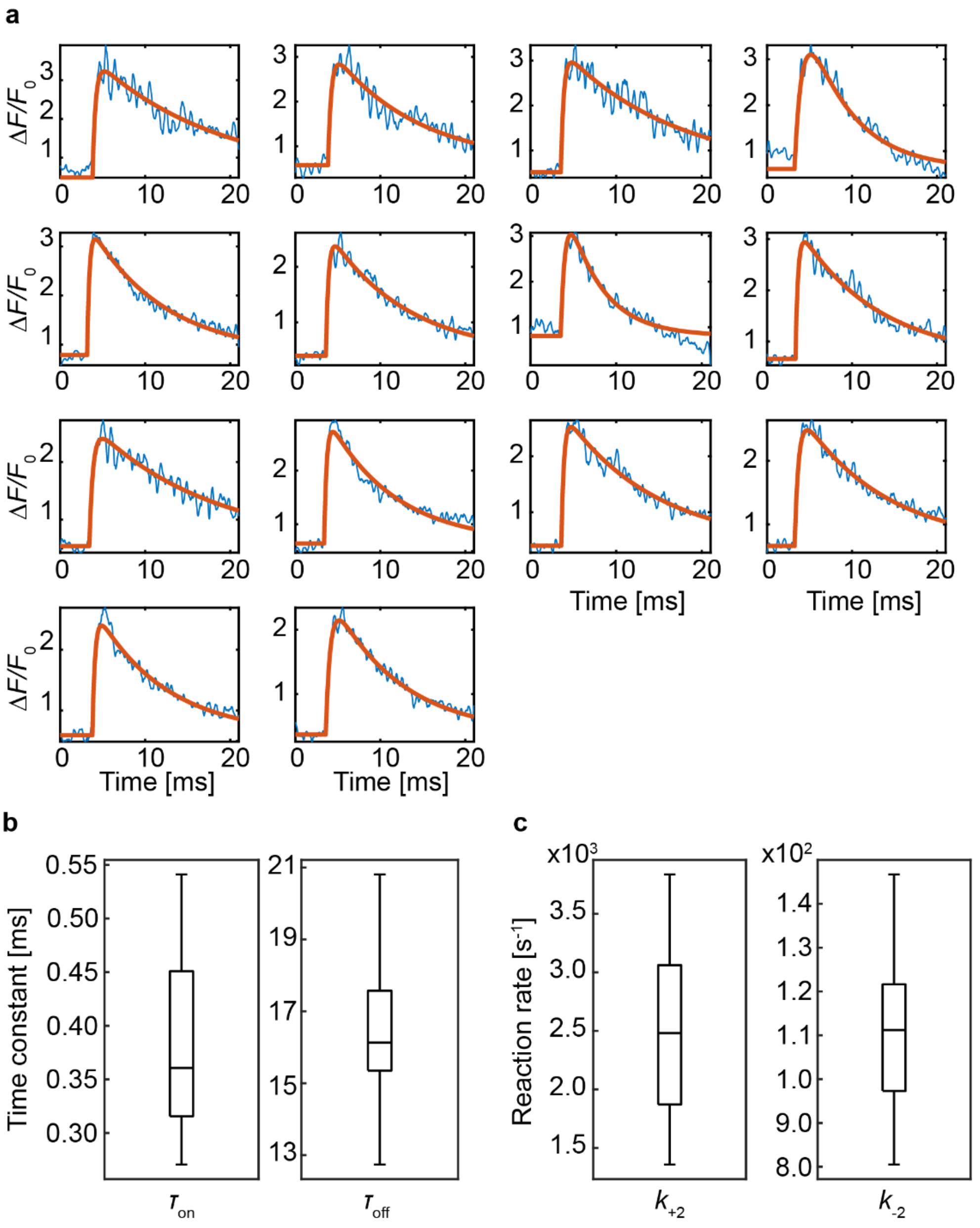


**Extended Data Figure 11: Extracting iGluSnFR kinetics from synaptic imaging data.**
**a)** Example traces of point scan (64 kHz sampling) of iGluSnFR signal in response to a single action potential. Blue lines: 14 success trials from 1 bouton. Red lines: double exponential fits. **b)** Median $\tau_{on}$ : 0.36 ms, Interquartile range IQR: 0.31-0.45 ms; median $\tau_{off}$ : 16.1 ms, IQR: 15.4-17.6 ms ($n$ = 3 boutons, 10 to 20 trials). **c)** Reaction rates, assuming two consecutive first-order reactions: $\text{Glu-iGluSnFR}_{dim} \xrightarrow{k_{+2}} \text{Glu-iGluSnFR}_{bright} \xrightarrow{k_{-2}} \text{Glu + iGluSnFR}_{dim}$
The simplified reaction scheme accounts for the fast diffusion of free glutamate out of the synaptic cleft (no equilibrium) and the slow conformational change dominating the kinetics of fluorescence increase ($k_{+2} >> k_{-1}$). Median values (median $k_{+2}$: 2481 $s^{-1}$, IQR: 1898-3020 $s^{-1}$, median $k_{-2}$: 111 $s^{-1}$, IQR: 97-122 $s^{-1}$) were used for the MCell model.
